# MDTraj: a modern, open library for the analysis of molecular dynamics trajectories

**DOI:** 10.1101/008896

**Authors:** Robert T. McGibbon, Kyle A. Beauchamp, Christian R. Schwantes, Lee-Ping Wang, Carlos X. Hernández, Matthew P. Herrigan, Thomas J. Lane, Jason M. Swails, Vijay S. Pande

**Author notes:** Electronic mail (Robert T. McGibbon).

## Abstract

*Summary:* MDTraj is a modern, lightweight and efficient software package for analyzing molecular dynamics simulations. MDTraj reads trajectory data from a wide variety of commonly used formats. It provides a large number of trajectory analysis capabilities including RMSD, DSSP secondary structure assignment and the extraction of common order parameters. The package has a strong focus on interoperability with the wider scientific Python ecosystem, bridging the gap between molecular dynamics data and the rapidly-growing collection of industry-standard statistical analysis and visualization tools in Python.

*Availability:* Package downloads, detailed examples and full documentation are available at http://mdtraj.org. The source code is distributed under the GNU Lesser General Public License at https://github.com/simtk/mdtraj.

## I. INTRODUCTION

Molecular dynamics (MD) simulations yield a great deal of information about the structure, dynamics and function of biological macromolecules by modeling the physical interactions between their atomic constituents. Modern MD simulations, often using distributed computing, graphics processing unit (GPU) acceleration, or specialized hardware can generate large datasets containing hundreds of gigabytes or more of trajectory data tracking the positions of a system’s atoms over time. In order to use these vast and information-rich datasets to understand biomolecular systems and generate scientific insight, further computation, analysis and visualization is required^1^.

Within the last decade, the Python language has become a major hub for scientific computing, with a wealth of high-quality open source packages, including those for interactive computing^2^, machine learning^3^ and visualization^4^. The environment is ideal for both rapid development and high performance, as computational kernels can be implemented in C and FORTRAN but available within a user-friendly environment.

In the MD community, the benefits of integration with such industry standard tools has not yet been fully realized because of a tradition of custom file formats and command-line analysis^5, 6^. In order to bridge this gap, we have developed MDTraj, a modern, open and lightweight Python library for analysis and manipulation of MD trajectories with the following goals:

1. To serve as a *bridge* between MD data and the modern statistical analysis and scientific visualization software ecosystem in Python.
2. To support a wide set of MD data formats and computations.
3. To run extremely rapidly on modern hardware with efficient memory utilization, enabling the analysis of large datasets.

## II. CAPABILITIES AND IMPLEMENTATION

*Wide range of data formats:* MDTraj can read and write from a wide range of data formats in use within the MD community, including `RCSB pdb, GROMACS xtc and trr, CHARMM / NAMD / OpenMM dcd, TINKER arc, AMBER NetCDF, binpos, mdcrd` and `prmtop` files. This wide support enables consistent interfaces and reproducible analyses regardless of users’ preferred MD simulation packages.

*Easy featurization:* Many data-analysis methods for MD involve either (a) extracting a vector of order parameters of each simulation snapshot or (b) defining a distance metric between snapshots. This category includes dimensionality reduction techniques such as principal components analysis (PCA) for constructing free-energy landscapes, as well probabilistic models like Markov state models.

MDTraj makes it very easy to rapidly extract these representations. It includes an extremely fast minimal root mean squared deviation (RMSD) engine capable of operating near the machine floating point limit described by^7^. Functions for DSSP secondary-structure assignment^8^, solvent accessible surface area determination and the extraction of internal degrees of freedom are similarly optimized in C with extensive use of vectorized intrinsics.

~~~
**import** mdtraj as md
t = md. load (‘trajectory. pdb’)
**from** itertools **import** combinations
pairs = combinations (**range** (t. n_atoms), 2)
X = md. compute_distances (t, pairs)

**import** matplotlib. pyplot as plt
**from** sklearn. decomposition **import** PCA
pca = PCA(n_components=2)
Y = pca.fit_transform (X)
plt. hexbin (Y[:, 0], Y[:, 1], bins=‘log’)
~~~

*Interactive visualization:* These fast computational routines make MDTraj ideal for interactive calculation and exploratory analysis, using the extensive machine learning, statistics and visualization packages in the scientific python community. Furthermore, MDTraj includes an interactive WebGL 3D protein viewer in the IPython notebook based on iview^9^, shown in Fig. 2.

**Figure 1:**
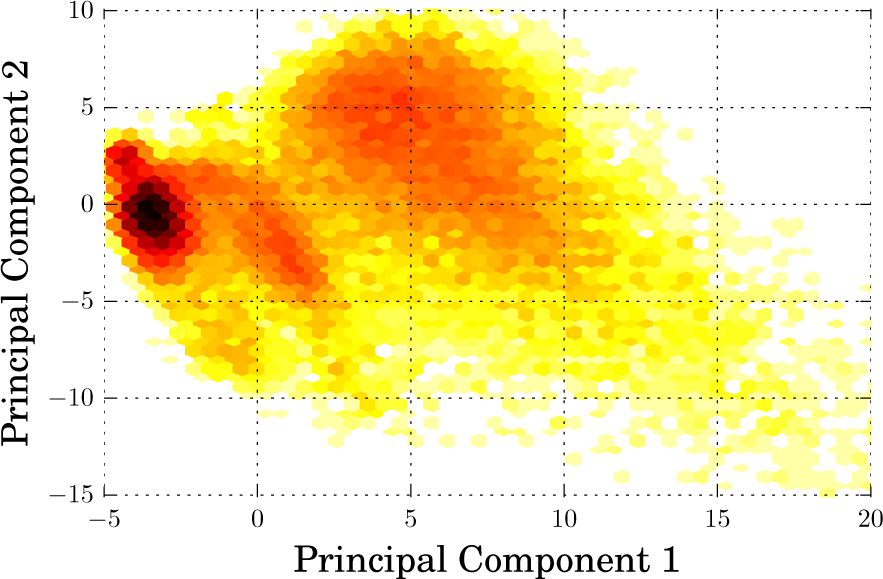
Demonstration of principal components analysis (PCA) with MDTraj, scikit-learn and matplotlib.

**Figure 2:**
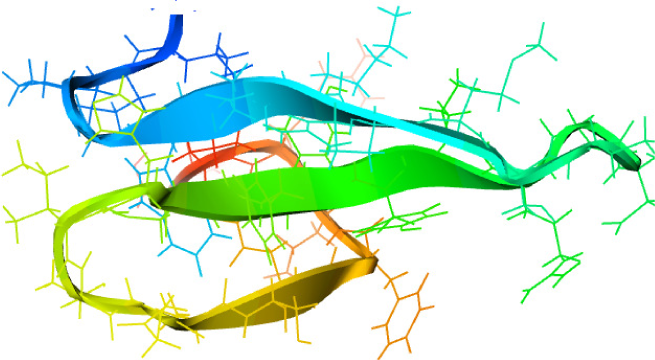
MDTraj’s WebGL-based protein and trajectory viewer.

The capabilities of MDTraj serve as a *bridge*, connecting MD data with statistics and graphics libraries developed for general data science audiences. The key advantage of this design, for users, is access to a much wider range of state-of-the-art analysis capabilities characterized by large feature sets, extensive documentation and active user communities.

A demonstration of this integrative workflow is shown in Fig. 1, which combines MDTraj with the scikit-learn package for PCA and matplotlib for visualization, to determine high-variance collective motions in a protein system. While PCA is a widely used method that is included in a variety of MD analysis packages, the advantage of integrating with the wider data science community is immediately evident when moving on to more complex statistical analysis. For example, a variety of sparse and kernelized PCA-like methods have been recently introduced in the machine learning community, and may be quite powerful for analyzing more complex protein systems. Because of its open and interoperable design, these cutting-edge statistical tools are readily available to MD researchers with MDTraj, without duplication of developer efforts and independent of the particular MD software used to perform the simulations.

## III. TESTING AND DEVELOPMENT

The development and engineering of MDTraj incorporates modern best practices for scientific computing^10^, and contains over 900 tests for individual components. These tests are continually run on each incremental contribution on both Windows and Linux platforms, using multiple versions of Python and the required libraries. The project is licensed under the GNU Lesser General Public License, and its design and development takes place openly on Github at https://github.com/simtk/mdtraj. More information is available at http://mdtraj.org.

*Funding:* National Institutes of Health (R01-GM62868, P30-CA008748); National Science Foundation (MCB-0954714). *Conflicts of Interest:* None declared.

